# Genome Architecture Shapes Transcriptional Responses to DNA Supercoiling in a Multicellular Organism

**DOI:** 10.1101/2025.01.03.631213

**Authors:** Ana Karina Morao, Almira Chervova, Ana Kotte, Yuya Zhao, Ivano Legnini, Sevinc Ercan, Germano Cecere

## Abstract

DNA supercoiling is an intrinsic consequence of transcription that must be resolved to maintain proper gene expression. How DNA supercoiling shapes transcription dynamics in chromatinized animal genomes remains unclear. Here, we acutely depleted topoisomerases I and II in *Caenorhabditis elegans* and applied nascent transcription profiling, nuclear and total RNA-seq, histone modification mapping, and long-read sequencing to capture the immediate transcriptional and chromatin responses to topological stress. We show that the genomic context influences the effect of supercoiling on transcription initiation, elongation and coordinated expression of adjacent genes. The impact of supercoiling on transcription initiation is not uniformly repressive but instead depends on the orientation and proximity of neighboring genes. We find that negative supercoiling promotes coordinated expression of divergent gene pairs, while positive supercoiling inhibits and uncouples expression of convergent genes. DNA supercoiling hinders transcription elongation globally resulting in reduced production of longer transcripts and overall shortening of poly(A) tails. While DNA supercoiling generated by neighboring transcription impacts initiation, elongation defects are driven by local supercoiling generated by the gene’s own transcription. These elongation effects are not accompanied by global changes in elongation-associated histone modifications but coincide with modest reductions in promoter and enhancer marks. Our findings reveal a directional mechanism by which genome architecture shapes transcriptional responses to DNA supercoiling, uncovering a DNA topological basis for coordinated gene expression in a multicellular organism.

## Introduction

DNA-templated processes including transcription and replication involve the opening of the double helix and the translocation of polymerases that force DNA to rotate around its axis. This rotation results in positively supercoiled DNA ahead of the polymerase and negatively supercoiled DNA behind^1^. While negative supercoiling contribute to strand separation and to the binding of the transcription machinery and positive supercoiling helps destabilizing nucleosomes for transcription progression^2–7^, excessive or unresolved torsional stress is detrimental to transcription and must be relieved to maintain proper gene expression^8–10^.

Topoisomerases play a central role in controlling DNA supercoiling. Type I topoisomerases cleave and re-ligate a single DNA strand to resolve supercoiling, whereas Type II topoisomerases cleave both DNA strands to resolve supercoiling, DNA catenates, and knots^11^. Genetic and biochemical studies in multiple organisms have shown that both enzymes can support transcription, with partially redundant roles^9,12–15^. However, most in vivo studies rely on steady-state RNA measurements or RNA Pol II occupancy data, which do not discern direct from indirect effects, nor distinguish whether topoisomerases primarily affect transcription initiation, elongation, or termination. Direct measurements of nascent transcription are therefore critical to dissect the role of topoisomerases at the different stages of transcription.

Beyond single genes, supercoiling can propagate along the chromatin fiber and influence nearby transcription. In yeast, negative supercoiling generated by transcription has been shown to diffuse up to 1.5 kb upstream of transcription start sites (TSS)^16^, raising the possibility that neighboring genes might influence each other’s expression through propagation of torsional stress. Computational models and bacterial studies support this view, suggesting that the orientation of neighboring genes determines the direction and outcome of supercoiling: divergent genes, transcribing away from each other, promote mutual activation via negative supercoiling, whereas convergent genes generate positive supercoils, which are repressive^17–24^. Whether similar architecture-dependent effects occur in eukaryotic animal genomes, where genomic distances are larger and multiple mechanisms regulate genome compaction, remains unclear.

Here, we combined acute auxin-inducible depletion of topoisomerases I and II in *Caenorhabditis elegans* with genome-wide profiles of nascent transcription (GRO-seq), nuclear and total RNA abundance, poly(A) tails (FLAM-seq), and histone modifications (Cut&Tag). This integrative approach allowed us to investigate how DNA supercoiling impacts transcription initiation, elongation and termination, and reveal its role in the coordinated expression of adjacent genes. The effect of topoisomerase depletion on transcription initiation varied according to the orientation and proximity of neighboring genes: initiation was reduced across genes having close neighbors transcribing towards the gene, whereas genes with far away neighbors transcribing away from the gene displayed higher initiation. On the other hand, neighboring transcription had little effect on transcription elongation. Instead, the effect on elongation scaled with the length of the gene: elongation was strongly hindered across long genes that accumulate higher levels of DNA supercoiling compared to shorter genes. These elongation effects were not accompanied by global changes in elongation-associated histone modifications, instead we observe modest reductions in promoter and enhancer histone marks. In line with modelling predictions, we observe that negative supercoiling promotes coordinated expression of divergent gene pairs, while positive supercoiling uncouples expression of convergent genes. Our findings uncover a directional logic by which the physical organization of genes, including their orientation and proximity, shapes transcriptional responses to supercoiling. These results suggest that genome architecture encodes a regulatory layer in which DNA supercoils contribute to local gene coordination across a eukaryotic multicellular genome.

## Results

### Topoisomerase depletion alters transcription dynamics genome-wide

To examine the genome-wide impact of DNA supercoiling on transcription, we acutely depleted the two main topoisomerases in *C. elegans*, TOP-1 and TOP-2, and profiled nascent transcription using GRO-seq. By combining short-term topoisomerase depletion (1 hour) with a direct readout of transcriptional activity, we aimed to capture the immediate effects of supercoiling accumulation on transcription.

To analyze the individual and combined contributions of TOP-1 and TOP-2, we performed GRO-seq in strains carrying *degron::GFP* of TOP-1, TOP-2, or both, along with a somatic TIR-1 transgene^25^ (Supplementary Figure 1A). As controls, we used the degron-tagged strains that were not treated with auxin (no-auxin control) as well as a strain without a degron tag but containing the TIR-1 transgene, which we treated with auxin (auxin control) (Supplementary Figure 1A). Treatment with auxin for one hour resulted in the rapid depletion of the GFP signal, confirming topoisomerase degradation (Supplementary Figure 1B).

In control animals, GRO-seq signal was distributed evenly across the gene bodies, with a modest peak at transcription start sites (TSS) and a pronounced peak near transcription end sites (TES), consistent with slow 3′-end processing in *C. elegans*^26,27^ (Figure 1A and Supplementary Figure 1C). Upon TOP-1 degradation, we observed a redistribution of GRO-seq signal, with some genes displaying an increase within the gene body and reduced signal at TESs (Figure 1A and Supplementary Figure 1C). In contrast, TOP-2 depletion alone caused no significant changes (Figure 1A and Supplementary Figure 1C), consistent with a more dominant role for TOP-1 in resolving transcription-induced supercoiling ^6,13,28^.

**Figure 1.**
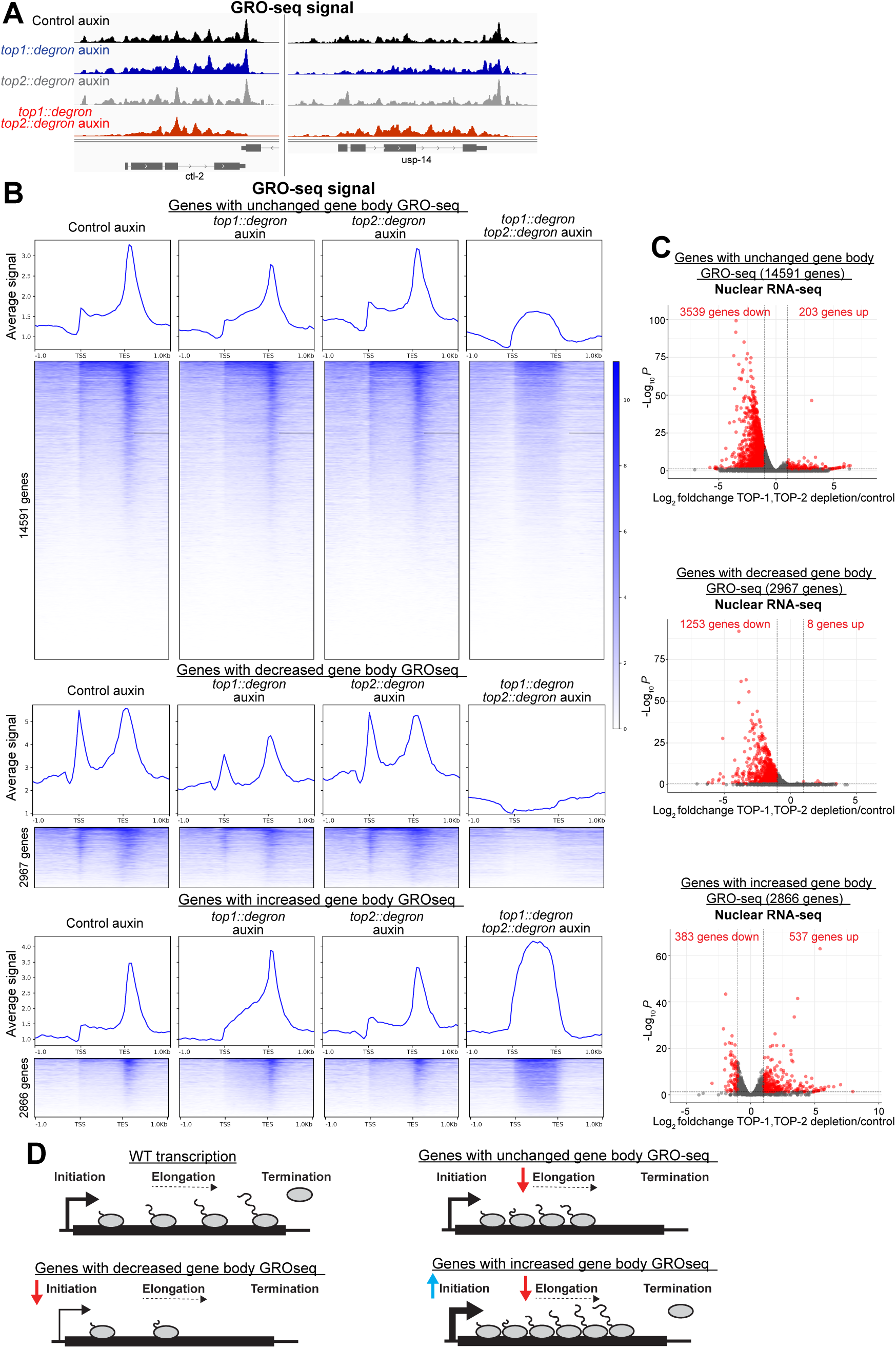
Topoisomerase depletion impacts transcription dynamics genome-wide. **A** Genome browser (IGV) view of GRO-seq profiles in control auxin, TOP-1, TOP-2 and TOP-1/TOP-2 depleted conditions at representative genes. **B** Heatmaps showing GRO-seq signal across genes with unchanged, decreased and increased levels of transcribing RNA Pol-II within the gene body. GRO-seq signal is shown for control auxin, *top-1::degron, top-2::degron and top-1::degron,top-2::degron* auxin conditions. Average plots are included at the top of the heatmaps. **C** Volcano plots depicting nuclear RNA-seq changes across genes with unchanged, decreased and increased levels of GRO-seq signal along the gene body. **D** Model of transcription steps affected by topoisomerase depletion in genes with unchanged, reduced and increased gene-body GRO-seq signal.

Combined degradation of TOP1 and TOP2 lead to widespread disruption of transcriptional profiles, characterized by the loss of the TSS and TES peaks and accumulation of GRO-seq signal within the gene body (Figure 1A and Supplementary Figure 1C). Differential expression analysis using GRO-seq data identified genes with decreased (n=2967), increased (n=2866), or unchanged (n=14591) signal across gene bodies (Figure 1B). Genes with decreased signal showed nearly complete loss of RNA Pol II transcription across the entire gene, suggesting that excessive supercoiling impairs transcription initiation. In contrast, genes with increased or unchanged GRO-seq signal, displayed strong enrichment within the gene body and loss of TSS and TES peaks (Figure 1B), indicating that transcription elongation or pausing is perturbed, leading to accumulation of transcribing RNA Pol II along the gene body. This interpretation is supported by RNA Pol II ChIP-seq performed in the same conditions, which displayed similar changes (Supplementary Figure 1D). The differences in GRO-seq profiles indicate that the effect of topoisomerase depletion is not homogenous across all genes, with some genes showing almost complete loss of transcribing RNA Pol-II, and other genes displaying varying levels of RNA Pol-II accumulation within the gene body.

To assess how these GRO-seq profiles translate into RNA output, we performed nuclear total RNA-seq with spike-in controls to enable quantitative measurements. Among the genes with decreased GRO-seq signal, 42% were downregulated at the nuclear RNA level, consistent with impaired transcription, while only 0.3% were upregulated (Figure 1C-D). Genes with unchanged gene body GRO-seq signal showed predominantly reduced RNA abundance (24% downregulated), suggesting that RNA Pol II accumulation within gene bodies largely reflects hindered elongation, which reduces transcript output. Among genes with increased GRO-seq signal, 13% were downregulated and 19% upregulated, suggesting that impaired elongation can be offset by enhanced transcription initiation resulting in higher mRNA production for a subset of these genes.

Together, these results indicate that topoisomerase depletion alters transcription dynamics genome-wide, with gene-specific effects on initiation, elongation and 3’ RNA Pol II pausing.

### The effect of TOP-1 and TOP-2 depletion on transcription initiation is determined by gene orientation and distance from neighbors

We next focused on how topoisomerases depletion affects transcription initiation. Based on our GRO-seq data we identified genes showing reduced or increased levels of transcribing RNA Pol-II along the gene body. Given that 5’ RNA Pol II pausing is not widespread in *C. elegans*^26,27^, GRO-seq signal along the gene body largely reflects the combined output of transcription initiation, elongation and 3’ pausing. Thus, genes with increased GRO-seq signal likely experience higher transcription initiation rates and/or stronger elongation defects. Conversely reduced GRO-seq signal indicates impaired initiation.

To identify features influencing these different outcomes, we examined genomic parameters known to impact the supercoiling state of a gene. First, we assessed intergenic distance, as supercoils can propagate and affect neighboring genes. Genes with decreased transcription initiation had significantly closer neighbors (median intergenic distance: 371 bp) than those with increased (1057 bp) or unchanged (861 bp) signal (Figure 2A), suggesting that topoisomerase activity is particularly important for transcriptional initiation in densely packed genomic regions.

**Figure 2.**
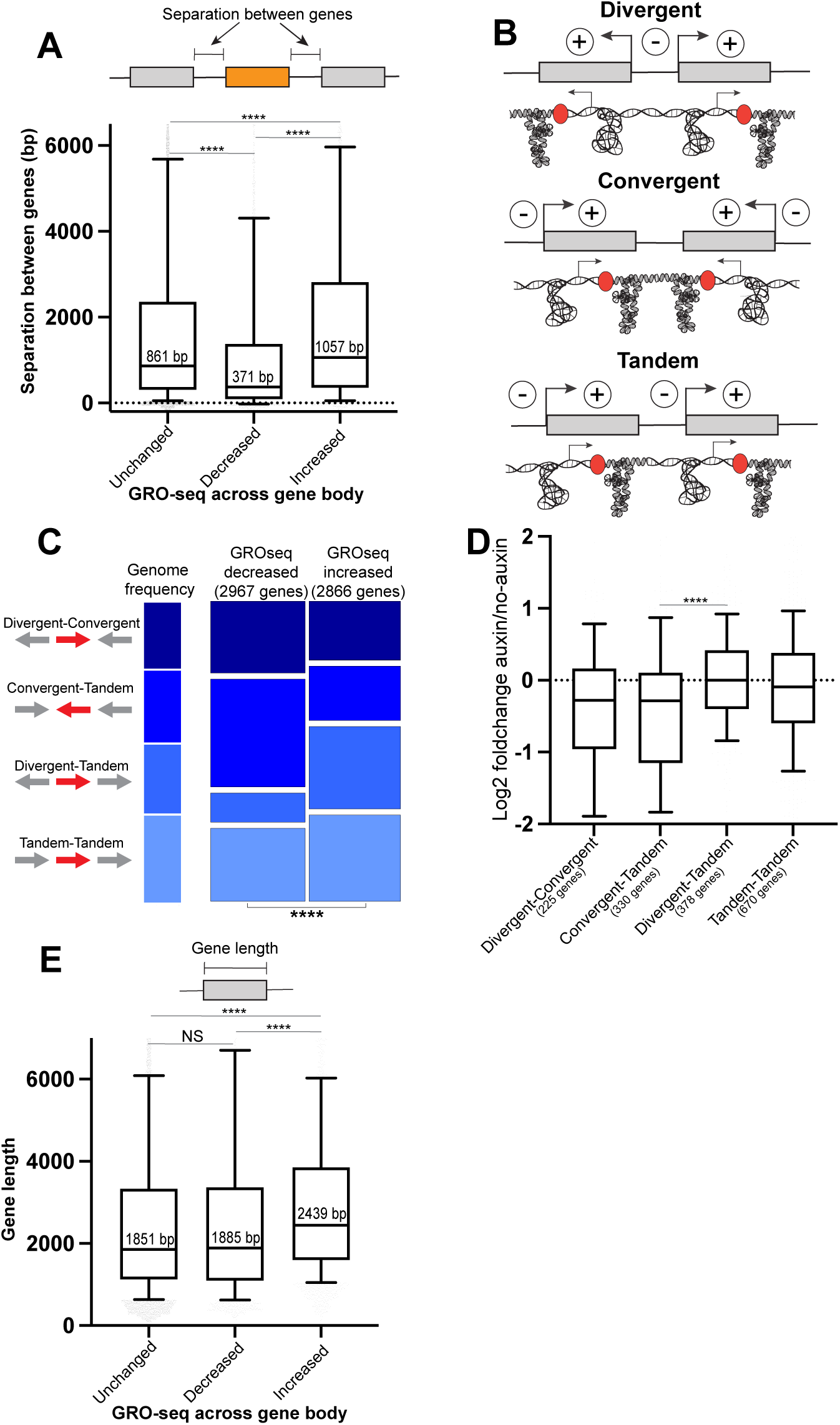
Genomic features determining the effect of TOP-1 and TOP-2 depletion on transcription initiation. **A** Distribution of the intergenic distances between genes neighbors is shown for genes with unchanged, decreased and increased levels of GRO-seq signal. Boxplots display median (line), first, and third quartiles (box), and 90th/10th percentile values (whiskers). Asterisks indicate significance of two-tailed p-values calculated using Mann-Whitney-Wilcoxon tests. **B** Schematic representation of gene pairs with different orientations. Negative and positive supercoiling generated by their transcription is indicated by (-) and (+) and represented by under and over-wounded DNA. **C** Genes were classified based on the orientation of their upstream and downstream neighbors into four groups colored in different tonalities of blue: Divergent-Convergent, Convergent-Tandem, Divergent-Tandem and Tandem-Tandem. The height of the of the bars represents the proportion of genes belonging to each orientation group. This proportion is shown for the genome (left) and for genes with increased and decreased GRO-seq signal (right). Asterisks at the bottom indicate significance of the difference between genes with decreased and increased GRO-seq (chi-square test p-value <10^-6^). **D** GRO-seq log2 fold changes between the auxin and no-auxin conditions are shown for Divergent-Convergent, Convergent-Tandem, Divergent-Tandem and Tandem-Tandem genes with neighbors within a distance between 100-600 bp. Boxplots display median (line), first, and third quartiles (box), and 90th/10th percentile values (whiskers). For comparison between Divergent-Tandem and Convergent-Tandem genes, two-tailed p-values were calculated using Mann-Whitney-Wilcoxon tests. The number of genes in each group is shown in parenthesis. **E** Distribution of gene length is shown for genes with unchanged, decreased and increased levels of GRO-seq signal. Boxplots display median (line), first, and third quartiles (box), and 90th/10th percentile values (whiskers). Asterisks indicate significance of two-tailed p-values calculated using Mann-Whitney-Wilcoxon tests.

Second, we analyzed gene orientation, which influences the type of DNA supercoiling that propagates towards a gene^8,29^. Neighbors in a divergent orientation generate negative supercoiling between each other, whereas those in a convergent orientation produce positive supercoiling (Figure 2B). Tandem gene pairs are predicted to have neutral or cancelling effects (Figure 2B). Given that negative and positive supercoiling impact transcription-related processes in different ways^8,29^, we hypothesized that accumulation of supercoiling upon topoisomerase depletion would have different outcomes for a gene depending on the orientation of its neighbors. To investigate this, we classified genes based on the orientation of their upstream and downstream neighbors into four categories (Figure 2C). 1) Tandem-Tandem: where both the gene behind and the gene ahead are in a tandem configuration (→→→, arrows represent gene orientation). 2) Divergent-Tandem: where one neighbor is divergent and the other is tandem (←→→). 3) Convergent-Tandem: where one neighbor is convergent and the other is tandem (→←←). 4) Divergent-Convergent: where one neighbor is divergent and the other is convergent (←→←).

The distribution of these orientation classes differed significantly between genes with increased and decreased GRO-seq signal (chi-square test, p < 10⁻⁶ Figure 2C). Genes with increased GRO-seq signal upon TOP-1/TOP-2 depletion are enriched for genes with upstream and downstream neighbors transcribing away from the gene and therefore producing a negative supercoiling domain around the gene (Divergent-Tandem neighbors) (Figure 2C). The opposite is observed for genes with decreased GRO-seq signal, which are enriched for those with upstream and downstream neighbors transcribing towards the gene and thus producing positive supercoiling in the direction of the gene (Convergent-Tandem neighbors) (Figure 2C). Genes with neighboring genes transcribing both away from and towards the gene (Divergent-Convergent and Tandem-Tandem) did not show a strong difference in GRO-seq signal upon TOP-1/TOP-2 depletion, suggesting that the effect of opposing supercoils may cancel each other. Analysis of transcription based on nuclear RNA-seq was consistent with results using GRO-seq (Supplementary Figure 2A).

Of note, we observe that the separation distance between convergent genes is shorter than between divergent or tandem genes genome-wide (Supplementary Figure 2B). Since intergenic distance may confound orientation effects, we next compared GRO-seq signal among orientation classes within a matched range of intergenic distances (100–600 bp) (Supplementary Figure 2C). Even within this range, Convergent-Tandem genes showed the strongest transcriptional repression, reinforcing the inhibitory role of unresolved positive supercoiling on transcription initiation (Figure 2D).

Lastly, we considered gene length, as longer genes accumulate higher levels of DNA supercoiling as RNA Pol II translocate across longer distances.^1^ Gene size was not significantly different between genes with decreased GRO-seq signal and genes with unchanged levels, suggesting that gene length is not a determinant of lower transcription initiation (Figure 2E). Instead, genes with increased GRO-seq signal were significantly longer than unchanged genes (Figure 2E), suggesting that elongation defects contribute to their RNA Pol II accumulation. This complicates interpretation for long genes, as both initiation and elongation may be affected, as discussed further below.

### Topoisomerase depletion affects transcription elongation in a gene length dependent manner

Our GRO-seq data showed that topoisomerase depletion leads to the accumulation of transcribing RNA Pol II across gene bodies (Figure 1B), which is consistent with a defect in transcription elongation. To further understand how the excess of supercoiling results in impaired elongation, we investigated the influence of gene length, neighbor gene’s orientation, and separation distance on the RNA Pol II accumulation profile.

To explore the role of gene size on transcription elongation, we classified genes in different groups based on their length and plotted their average GRO-seq signal (Figure 3A). On short genes, depletion of topoisomerases resulted in increased GRO-seq signal across the gene body with a 3’ bias, suggesting that a fraction of the RNA Pol II that initiated was able to reach the end of the gene (Figure 3A). This 3’ bias becomes less pronounced as the gene size increases, with the longest genes showing a 5’ bias, indicating that most of the transcribing RNA Pol II stop before reaching the end of the gene (Figure 3A).

**Figure 3.**
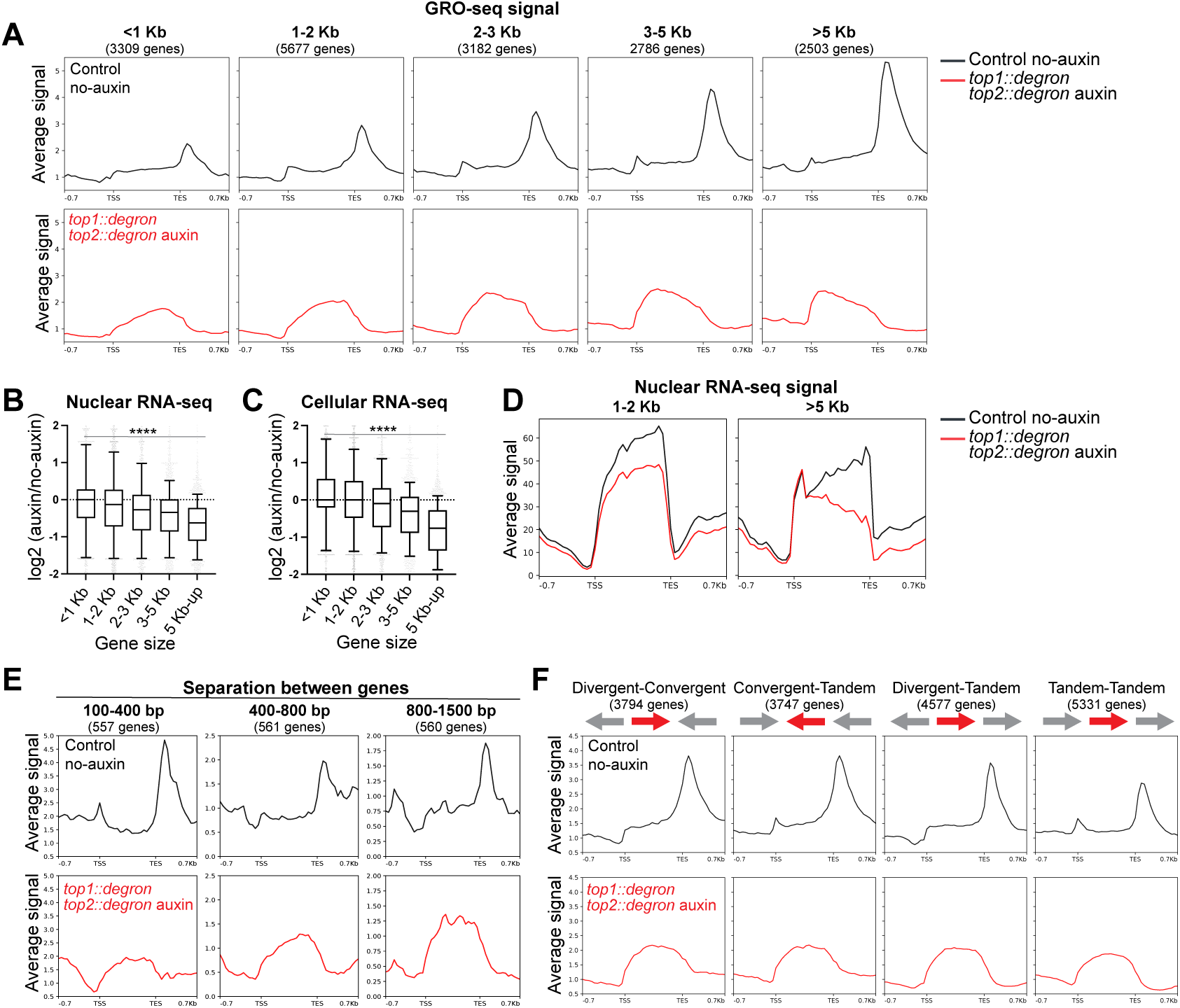
The effect of topoisomerase depletion on transcription elongation is determined by the amount of supercoiling generated within the gene. **A** Average GRO-seq scores in control (top, black) and TOP-1/TOP-2 depletion (bottom, red) are plotted across genes grouped based on their length into five categories: < 1 Kb, 1-2 Kb, 2-3 Kb, 3-5 Kb and > 5 Kb. The number of genes in each group is indicated in parenthesis. **B** Nuclear RNA-seq log2 fold changes between the auxin and no-auxin conditions are shown for genes of different sizes. Boxplots display median (line), first, and third quartiles (box), and 90th/10th percentile values (whiskers). For comparison between shortest (< 1Kb) and longest genes (> 5 Kb), two-tailed p-values were calculated using Mann-Whitney-Wilcoxon tests. **C** Cellular RNA-seq log2 fold changes between the auxin and no-auxin conditions are shown for genes of different sizes. For comparison between shortest (< 1Kb) and longest genes (> 5 Kb), two-tailed p-values were calculated using Mann-Whitney-Wilcoxon tests. **D** Average nuclear RNA-seq scores in control (black) and TOP-1/TOP-2 depletion (red) are plotted across short (1-2 Kb) and long genes (> 5 Kb). **E** Average GRO-seq scores in control (top, black) and TOP-1/TOP-2 depletion (bottom, red) are plotted across genes grouped based on the distance separating them from their adjacent neighbors (100-400 bp, 400-800 bp, 800-1500 bp). **F** Average GRO-seq scores in control (top, black) and TOP-1/TOP-2 depletion (bottom, red) are plotted across genes grouped based on the orientation of their upstream and downstream neighbors.

Consistent with the interpretation that a strong 5’ bias in the GRO-seq signal reflects hindered elongation, nuclear RNA-seq revealed that mRNA abundance decreased with increasing gene length (Figure 3B). In addition, nuclear RNA distribution along the gene mirrored the GRO-seq profile observed upon topoisomerase depletion, with long genes showing reduced transcript abundance towards the 3’ end, whereas short genes are evenly covered (Figure 3D). This suggests that long transcripts remain unfinished, whereas short transcripts are fully processed. To test this interpretation, we sequenced total cellular RNAs and indeed found that the transcript level of longer genes is reduced upon topoisomerase depletion (Figure 3C). Furthermore, TOP-1 degradation alone was sufficient to cause elongation defects in long genes, whereas TOP-2 depletion had no measurable effect (Supplementary Figure 3). Thus, TOP-1 is the primary topoisomerase supporting transcription elongation in *C. elegans*.

To assess whether neighboring transcription also influences elongation, we examined GRO-seq profiles of genes with different neighbor distances and orientations. No detectable differences were observed in the GRO-seq profile based on neighbor proximity (Figure 3E and Supplementary Figure 3C) or orientation (Figure 3F and Supplementary Figure 3D), suggesting that elongation defects are driven by supercoiling generated within the gene itself, rather than by neighboring transcription. Thus, unresolved transcription-induced supercoiling impairs elongation in a gene length-dependent manner, while neighbor effects are more important for transcription initiation.

In summary, analysis of transcription by GRO-seq, and the product of transcription using nuclear and cellular RNA-seq indicate that TOP-1/TOP-2 depletion have a significant impact on transcription elongation and completion that scales with gene length. Incomplete transcription resulting from unresolved transcription-mediated DNA supercoiling is consistent with previous work in yeast and mammals showing that topoisomerases are required for the expression of long genes ^14,15^.

### Topoisomerase depletion results in shorter poly(A) tails

Our GRO-seq data revealed a loss of the characteristic RNA Pol II peak near the transcription end site upon topoisomerase depletion (Figure 1B), suggesting a potential defect in 3’ mRNA processing. To test this, we performed full-length poly(A) and mRNA sequencing (FLAM-seq), which enables simultaneous measurement of poly(A) tail length and precise mapping of transcript 3’ends^30,31^.

We sequenced both nuclear and cellular mRNA pools. Poly(A) tails were longer in nuclear RNAs than in cellular RNAs (Figure 4A), consistent with previous observations^30^. Topoisomerase depletion did not alter the position of 3’ cleavage sites (Supplementary Figure 4A-B), indicating no widespread shift in polyadenylation sites usage. However, it led to a reduction in poly (A) tail length in both nuclear and cellular RNA transcripts (Figure 4A-B).

**Figure 4.**
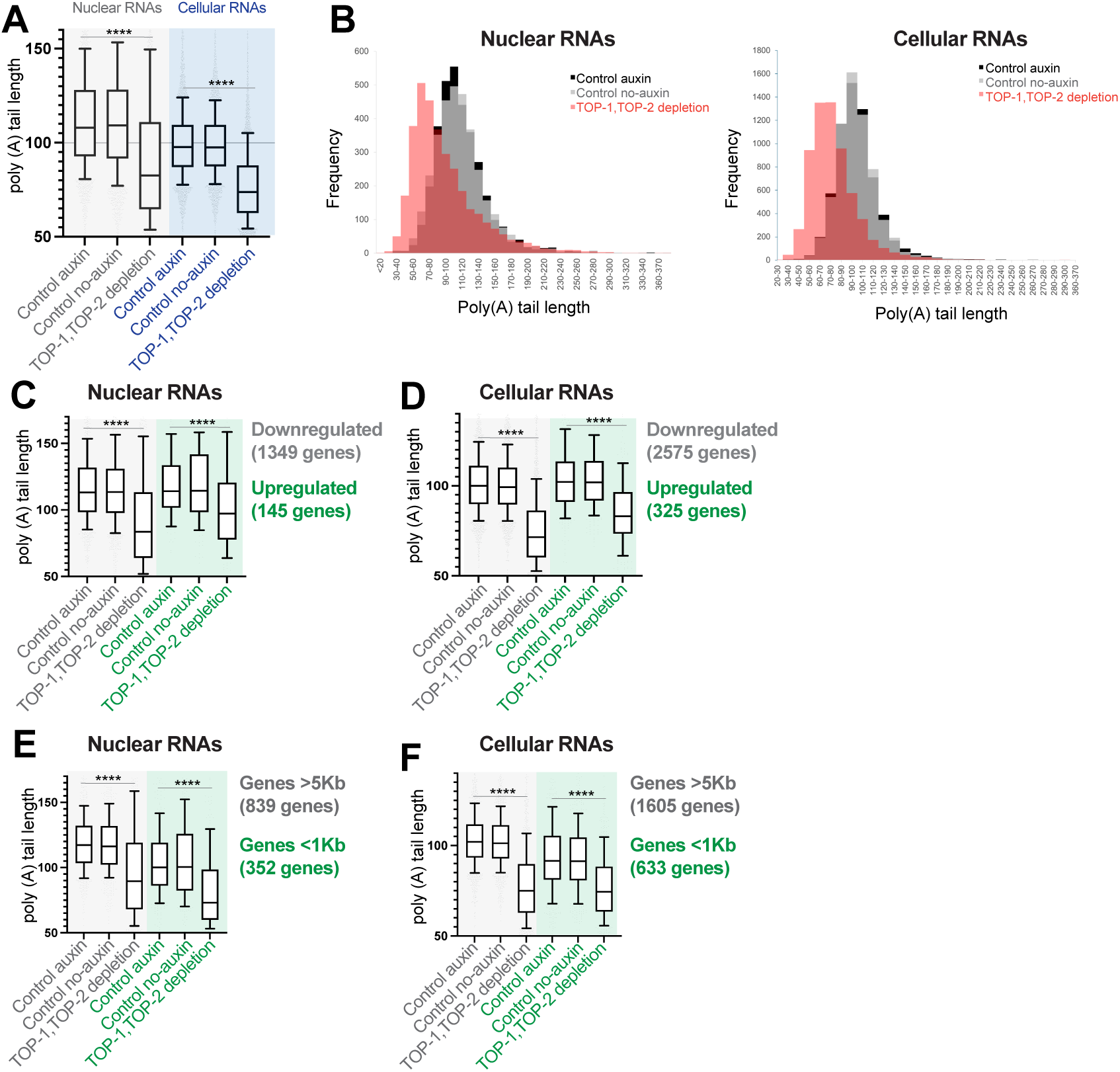
Topoisomerase depletion results in shorter poly(A) tails. **A** Boxplot of the distribution of poly(A) tail lengths for nuclear and cellular mRNAs in the controls and TOP-1/TOP-2 depletion. Boxplots display median (line), first, and third quartiles (box), and 90th/10th percentile values (whiskers). Asterisks indicate significance of two-tailed p-values calculated using Mann-Whitney-Wilcoxon tests. **B** Histograms of the distribution of poly(A) tail lengths of nuclear (left) and cellular (right) mRNAs in the controls and TOP-1/TOP-2 depletion. **C** Boxplot of the distribution of poly(A) tail lengths of nuclear mRNAs in the controls and TOP-1/TOP-2 depletion for genes found differentially expressed by nuclear RNA-seq upon topoisomerase depletion. Asterisks in box plots indicate significance of two-tailed p-values calculated using Mann-Whitney-Wilcoxon tests. **D** Boxplot of the distribution of poly(A) tail lengths of cellular mRNAs in the controls and TOP-1/TOP-2 depletion for genes found differentially expressed by nuclear RNA-seq upon topoisomerase depletion. **E** Boxplot of the distribution of poly(A) tail lengths of nuclear mRNAs in the controls and TOP-1/TOP-2 depletion for short (<1Kb) and long (>5Kb) genes. **F** Boxplot of the distribution of poly(A) tail lengths of cellular mRNAs in the controls and TOP-1/TOP-2 depletion for short (<1Kb) and long (>5Kb) genes.

One possibility is that shorter poly(A) tails could reflect a defect in polyadenylation itself, or alternatively, a secondary consequence of reduced mRNA production. In this scenario, older transcripts may undergo prolonged deadenylation in both the nucleus and cytoplasm^32^, leading to shorter average tail lengths.

To distinguish between these possibilities, we examined poly(A) tail lengths in transcripts that were upregulated upon topoisomerase depletion, reasoning that if reduced production were the main driver, upregulated genes should be less affected. Surprisingly, even upregulated transcripts exhibited shorter poly(A) tails in both nuclear and cellular fractions (Figure 4C-D, Supplementary Figure 4C-D), indicating a general effect on poly(A) tail length. However, the shortening was more pronounced in downregulated transcripts, suggesting that the rate of mRNA production plays a role in the level of deadenylation (Figure 4C-D, Supplementary Figure 4C-D).

A similar trend was observed with gene length. Longer genes, which show stronger elongation defects and lower transcript levels, exhibited greater poly(A) tail shortening in the cellular RNA fraction than short genes (Figure 4E-F, Supplementary Figure 4E-F). These findings suggest that altered transcription dynamics upon topoisomerase depletion are accompanied by genome-wide shortening of poly(A) tails, with the strongest effect on genes with reduced expression.

### Topoisomerases shape promoter chromatin but do not globally alter elongation-associated histone marks

Transcription-generated torsional stress can affect nucleosome stability^6,33^, yet its impact on histone modification landscapes remains unexplored. Given the defects in transcription initiation and elongation observed upon topoisomerases depletion, we profiled the active elongation mark H3K36me3, and the promoter/enhancer marks H3K4me3 and H3K27ac using Cut&Tag. We also profiled the repressive modification H3K27me3.

Despite impaired transcriptional elongation, we observe no major changes in H3K36me3 levels, even across long genes where the transcriptional defects were strongest (Figure 5A, Supplementary Figure 5A). In contrast, H3K4me3 and H3K27ac showed both gains and losses that correlated with transcriptional changes. Notably, even genes with unchanged expression displayed a modest decrease in these marks around transcription start sites (Figure 5C-D), suggesting a local chromatin response to topoisomerase loss. Previous studies have suggested that topoisomerases interact with histone modifiers^34,35^. Consistent with this, ChIP-seq data show that TOP-1 and TOP-2^25^ bind to promoters and their occupancy positively correlates with H3K27Ac and H3K4me3 levels (Supplementary Figure 5B). Thus, the observed decrease in promoter-associated histone modifications may result either from loss of topoisomerase binding or from supercoiling-induced disruption of chromatin state.

**Figure 5.**
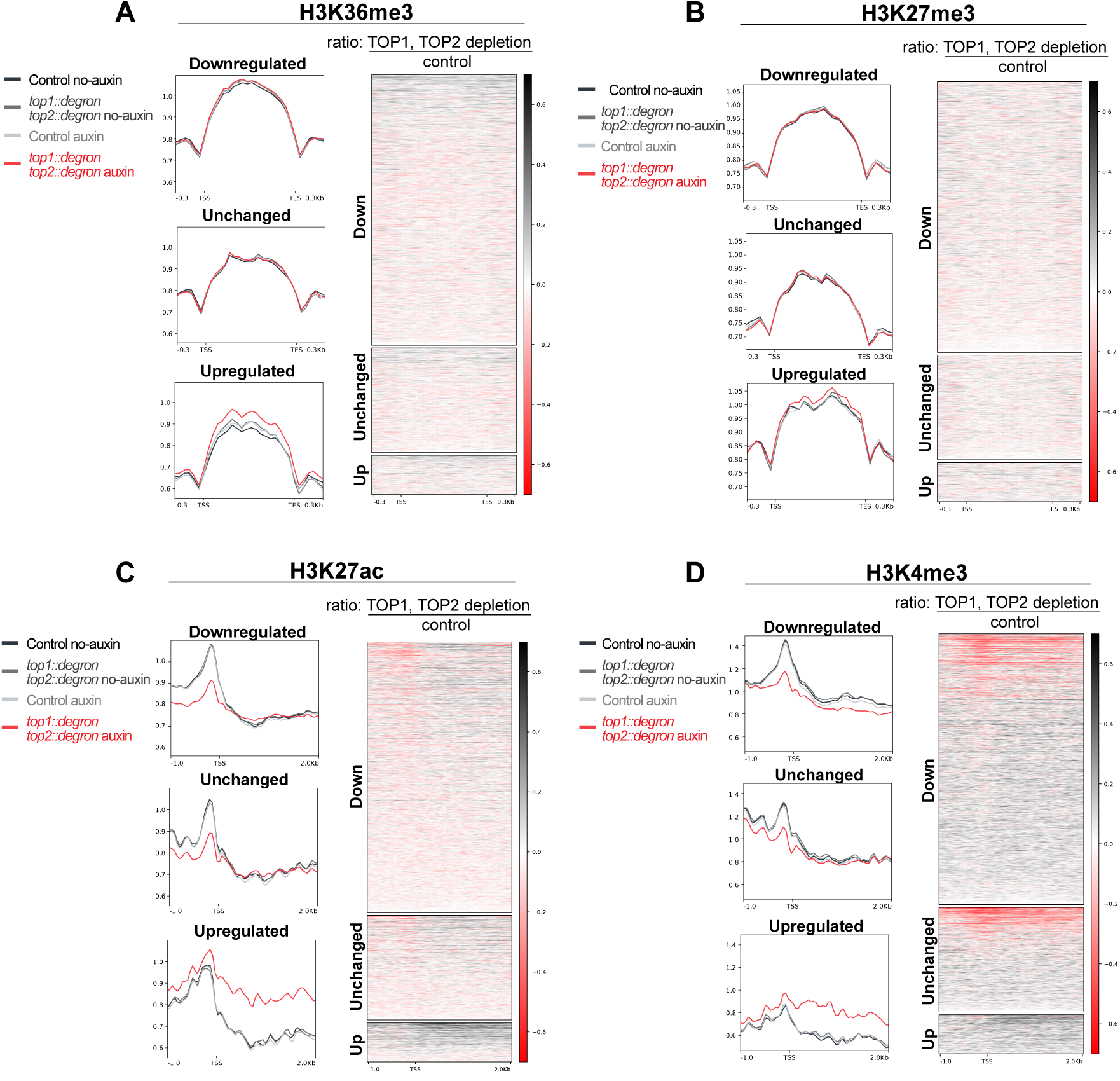
Impact of topoisomerases on histone modifications Right: Heatmap showing H3K36me3 **(A)**, H3K27me3 **(B)**, H3K27ac **(C)** and H3K4me3 **(D)** Cut & Tag ratios between TOP-1/TOP-2 depletion and control, across genes found differentially expressed by nuclear RNA-seq. **Left:** Average plots of the Cut & Tag signal including the no-auxin and auxin controls. For H3K27Ac and H3K4me3, signal around the TSS is shown. For H3K36 and H3K27me3, gene bodies were scaled to 1Kb. Unchanged genes include 2000 genes randomly selected for representation.

### Divergent genes exhibit coordinated transcription via negative supercoiling

We next asked whether transcription-induced supercoiling could influence the coordination of gene expression across neighboring genes. In bacteria, negative supercoiling between divergent genes enhances transcription, whereas positive supercoiling generated by convergent orientation inhibits transcription. Modelling studies have proposed similar architecture-dependent coupling in eukaryotes^17,18^.

To test this, we computed the correlation in transcriptional changes between divergent, convergent and tandem gene pairs, following TOP-1/TOP-2 depletion (Figure 6A). For this pairwise analysis, orientation was determined based on the closest neighbor. We found that divergent gene pairs showed a positive correlation, while convergent pairs showed no correlation, and tandem pairs showed a modest positive trend (Figure 6A). Therefore, we conclude that when DNA supercoiling levels are perturbed, divergent and to a lesser extent tandem gene pairs are affected synchronously while convergent genes have uncoordinated changes. These results suggest that negative supercoiling promotes co-regulation of divergent neighbors.

**Figure 6.**
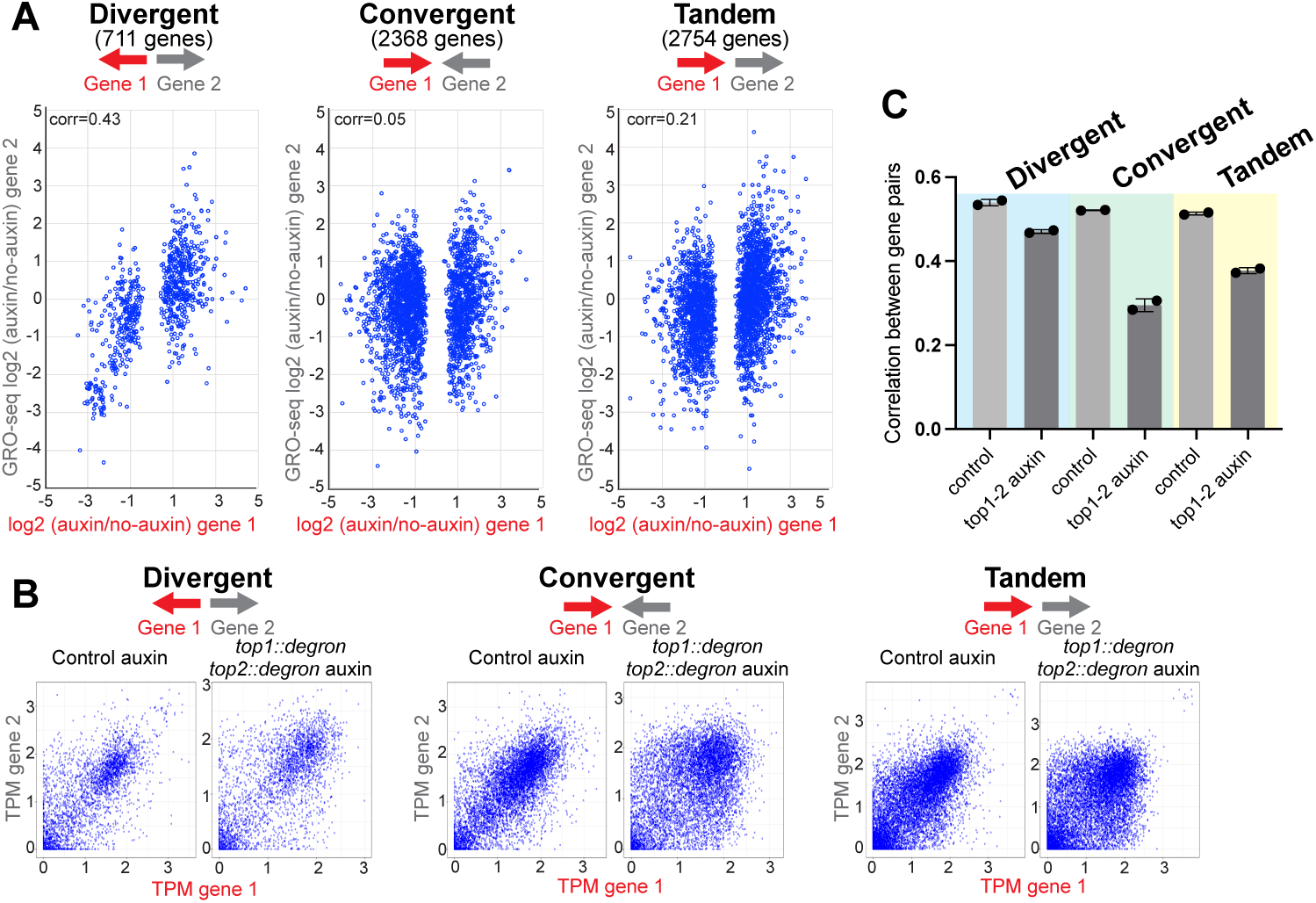
Divergent genes stimulate each other’s transcription through negative supercoiling. **A** Scatter plot showing GRO-seq log2-fold changes between the *top-1::degron,top-2::degron* auxin and no-auxin conditions for gene pairs with different orientations. **B** Scatter plot showing GRO-seq TPM values in control and *top-1::degron,top-2::degron* auxin for gene pairs with different orientations. **C** Kendal correlations between expression (GRO-seq TPM values) of gene pairs with different orientations are shown for control and TOP-1/TOP-2 depletion. Each dot represents a biological replicate.

Positive supercoiling has been proposed to inhibit the transcription of convergent genes. In this case, higher transcription of a gene should inhibit transcription of its convergent neighbor, resulting in a negative correlation. However, we observed no correlation between how the expression of convergent gene pairs change upon topoisomerase depletion (Figure 6A). Thus, we next asked to what extend neighboring genes expression correlate with each other. We plotted the correlation between the expression levels of gene pairs with different orientations in the control and TOP-1/TOP-2 depleted conditions. In the control, expression of neighbor genes for all the orientations showed a positive correlation (Figure 6B) suggesting that in the presence of topoisomerases, convergent genes do not inhibit each other. In the absence of topoisomerases, the correlation between the expression of convergent genes is strongly reduced, whereas the correlation between the expression of divergent genes is less affected (Figure 6B-C and Supplementary figure 6). This supports a model in which excessive positive supercoiling between convergent genes reduces their co-expression, thus decoupling convergent transcription. In contrast, excessive negative supercoiling between divergent genes reinforces co-expression even under topological stress.

## Discussion

In this study, we combined acute topoisomerase depletion with genome-wide nascent transcription profiling, nuclear and total RNA-seq, poly(A) tail sequencing, and histone modification mapping to dissect the impact of transcription-induced DNA supercoiling in a chromatinized, multicellular genome. This allowed us to interpret the complex impact of topoisomerase activity on transcription initiation and elongation, and on the transcriptional coupling between neighboring genes. Our approach revealed that the consequences of supercoiling on the different stages of transcription are highly context-dependent, varying with gene length, orientation, and intergenic spacing. These findings advance our understanding of how the physical arrangement of genes can encode a layer of gene regulation via DNA supercoiling.

### Transcription initiation, and not elongation, is influenced by neighboring genes

The torsional state of a gene is influenced by DNA supercoiling generated by its own transcription as well as the transcription of its neighboring genes. Here, we reveal that these two sources of DNA supercoiling have specific effects on transcription initiation and elongation. We observe that the rate of transcription initiation is impacted by DNA supercoiling coming from neighboring transcription, with genes having close neighbors transcribing towards the gene showing an inhibitory effect, whereas genes with far away neighbors transcribing away from the gene displaying a stimulatory effect (Figure 7). This is in line with experimental studies showing the enhancing effect of divergent transcription on specific example genes, and fits predictions made by modeling in both bacteria and eukaryotes^17–21^.

**Figure 7.**
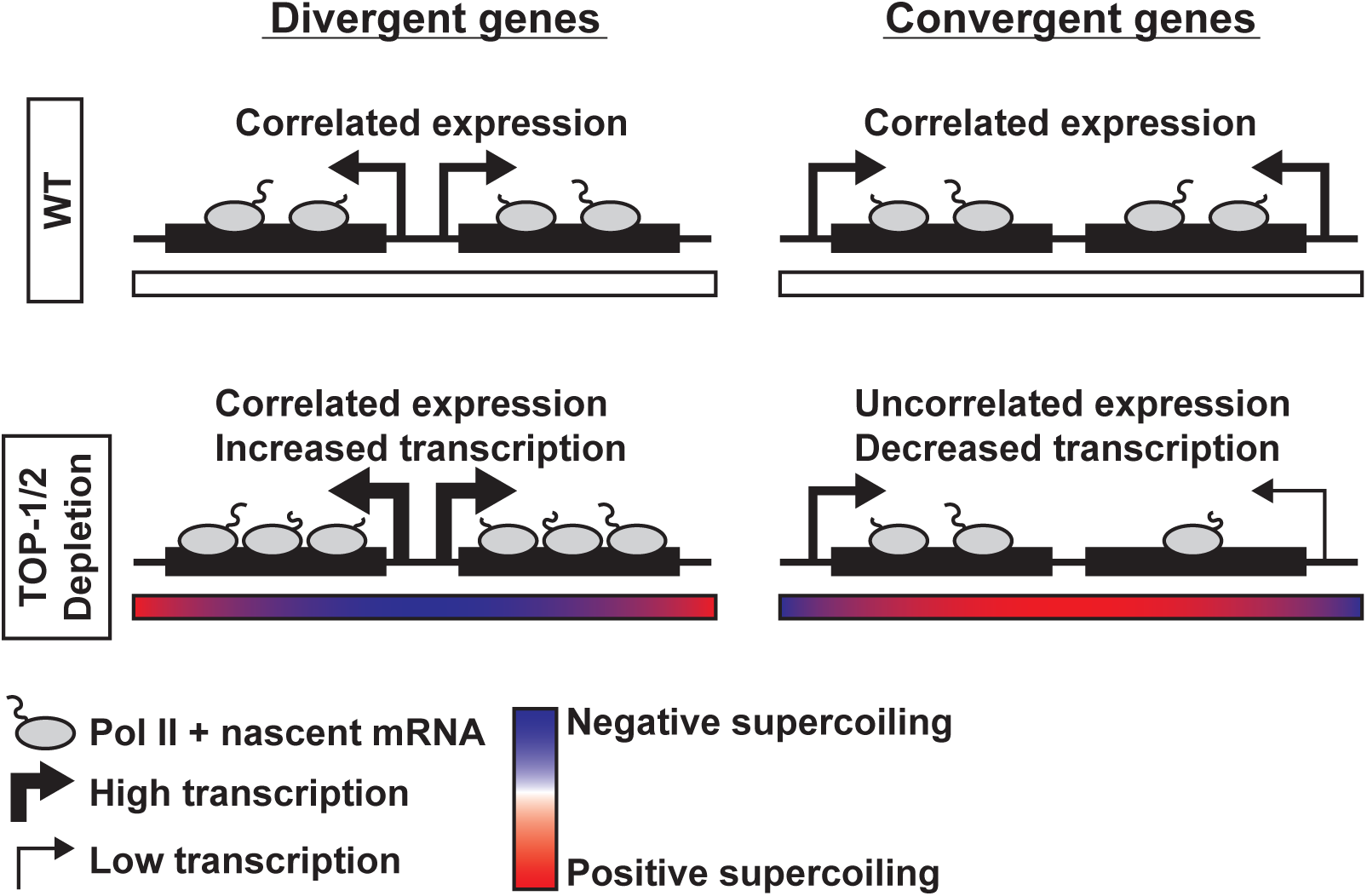
Model for the effect of topoisomerase depletion on the transcription of divergent and convergent genes.

Our results indicate that transcription elongation is influenced by DNA supercoiling originating in the gene itself. This is illustrated by the stronger inhibition of elongation observed in long compared to short genes, which differ in the amount of DNA supercoiling generated during their transcription. Neighboring transcription had no detectable influence on transcription elongation. On the other hand, transcription initiation was not differentially affected in short and long genes. Thus, while DNA supercoiling diffusing from neighboring transcription impacts initiation rate, torsional stress generated within the gene has no effect on initiation. Overall, these results support a model where topoisomerases activity at intergenic regions is required to support transcription initiation, while transcription elongation necessitates supercoiling relaxation within the gene.

### Negative and positive supercoiling have different effects on transcription

We observe that genes with upstream and downstream neighbors transcribing away from the gene (←→→, Divergent-Tandem genes) are less affected upon TOP-1/TOP-2 depletion, while genes with upstream and downstream neighbors transcribing towards the gene (→←←, Convergent-Tandem) become repressed. In the context of gene triplets, genes surrounded by divergent and tandem neighbors receive negative supercoiling from both sides, whereas genes flanked by convergent and tandem neighbors only receive positive supercoiling. Therefore, we propose that these two groups of genes are under the influence of opposite supercoiling states driving their differential response to topoisomerase depletion. In the case of genes with neighboring genes transcribed both away from and towards the gene (←→← Divergent-Convergent and →→→Tandem-Tandem), they receive both negative and positive supercoiling from their neighbors and show a milder response to topoisomerase degradation. We thus envision that for these genes, the effect of opposing supercoils may cancel each other. In summary our results are consistent with simulation studies predicting that negative supercoiling generated in the context of transcription of divergent genes acts to enhance their expression and positive supercoiling generated between convergent genes has an inhibitory effect^17,18,36^

In a recent study in yeast, topoisomerase depletion led to inhibition of genes in all orientations^37^, which led the authors to propose that the excess of both negative and positive supercoiling represses transcription by perturbing the binding of transcription factors. Negative supercoiling has been shown to promote binding of general transcription factors and other regulatory proteins^2–5,38^. Although we cannot exclude that excessive negative supercoiling resulting from the absence of topoisomerases could instead perturb protein binding to DNA, our results suggest that even in excess, negative supercoiling is less inhibitory to transcription than positive supercoiling.

### Coupling between convergent genes requires DNA supercoiling relaxation

Co-expression of adjacent genes is a strategy used by organisms to ensure coordinated expression of groups of genes sustaining transcriptional programs during development and in response to the environment ^39^. Diffusion of DNA supercoiling could act as a rapid mechanism to regulate expression of genes in close proximity ^17^. In the presence of topoisomerases, controlled levels of both positive and negative supercoiling might contribute to neighbors’ co-expression^40^ (Figure 7). In the absence of topoisomerases, excessive levels of negative and positive supercoiling have different effects on co-expression of neighbor genes with different orientations. While topoisomerase depletion did not affect correlated divergent genes, it perturbed coordinated expression of convergent genes (Figure 5B, (Figure 7)). These results suggest that coordinated expression of neighbor genes requires tight regulation of positive supercoiling levels, whereas negative supercoiling levels are more flexible. Thus, we propose that the divergent orientation affords a more robust configuration for coordinated expression.

### A non-catalytic role for topoisomerases in the regulation of histone modifications?

We hypothesized that DNA supercoiling could impact the deposition/removal of histone modifications by either its effect on transcription dynamics, which in turn would affect transcription-coupled histone modifications, or through its effect on nucleosome stability, which could impact how nucleosomes are displaced and reincorporated during transcription. We do not observe strong changes in the distribution of elongation-associated histone modification H3K36me3 upon topoisomerase depletion. This suggests that reduced transcription elongation in the absence of topoisomerases does not immediately perturb H3K36me3. Moreover, although topoisomerase inactivation was shown to result in higher nucleosome turnover within the gene body^6^, our data suggest that this does result in changes in the distribution of H3K36me3.

On the other hand, we observe a mild local reduction of H3K27ac and H3K4me3 around transcription start sites suggesting that DNA supercoiling could influence histone modifications associated with upstream regulatory elements. Another possibility is that this effect on H3K27ac and H3K4me3 is independent of the enzymatic activity of topoisomerases. A method developed to map catalytically engaged TOP-1 showed that it is depleted at promoters and enriched within the gene body^41^. Yet, chromatin immunoprecipitation experiments in various organisms showed TOP-1 and TOP-2 accumulation at promoters^25,28,41,42^. Therefore, binding at promoters is evolutionarily conserved but might not reflect catalytically engaged forms of topoisomerases. Instead, promoter associated topoisomerases might function through a non-catalytic mechanism, regulating transcription and chromatin structure through protein-protein interactions.

Taken together, our results reveal that the physical arrangement of genes within the genome, including their orientation and spacing, shapes the transcriptional response to supercoiling. By coupling genomic architecture with DNA mechanics, cells may encode a layer of transcriptional logic that supports local co-regulation, enhances robustness, or buffers topological stress. These findings suggest that DNA supercoiling may serve as an instructive signal, transmitting information across chromatin to coordinate gene expression.

## Acknowledgments

G.C., A.C. and A.K.M. have received funding from the Institut Pasteur and the CNRS. A.K.M. was also supported by a LabEx Revive fellowship ANR-10-LBX-73. S.E., Y.Z., and A.K.M and research in this article were supported by NIGMS of the National Institutes of Health under award number R35 GM130311. A.K. is a PhD student within the European School of Molecular Medicine (SEMM).

## Author contributions

A.K.M, G.C. and S.E. conceptualized the project and designed experiments. A.K.M. performed all experiments except for some ChIP-seq replicates and FLAM-seq. Y.Z. performed ChIP-seq experiments. A.K. and I.L. performed and analyzed FLAM-seq experiments. A.K.M. and A.C. performed bioinformatics analysis. A.K.M, G.C. and S.E. wrote the manuscript with contributions from the authors.

## Declaration of interests

The authors declare no competing interest.

## STAR methods

### Worm strains and growth

Worms were grown and maintained at 20°C on Nematode Growth Medium (NGM) plates containing the *E. coli* strain OP50. To isolate synchronized L2 worms, gravid adults were bleached in 0.5 M NaOH and 1.2% bleach, and embryos were hatched overnight in M9 buffer (22 mM KH2PO4 monobasic, 42.3 m in M Na2HPO4, 85.6 mM NaCl, 1mM MgSO4). Starved L1s were grown for 24 hours at 22°C in plates seeded with OP50 and the resulting L2 worms were used for auxin treatment. The *top-1::degron::GFP* and *top-2::degron::GFP* worms strains used in this study were previously described in Morao, *et al* 2022^25^

### Auxin treatment

A 400 mM auxin (indole-3-acetic-acid, Fisher 87-51-4) solution was prepared by resuspending auxin powder in 100% ethanol. 1 mM auxin plates were prepared by adding resuspended auxin to NGM media before pouring the plates. Around 2×10^5^ synchronized L2 worms obtained as described above were washed three times with M9 and divided in two aliquots. Half of the worms were transferred to NGM 15 cm plates supplemented with 1 mM of auxin. The other half was transferred to normal NGM 15 cm plates. Both auxin and no-auxin conditions were incubated for one hour at room temperature. Worms were then washed once with M9 and stored accordingly to future application. For GRO-seq and Cut & Tag, worms were stored in M9 buffer at-20 C till further processing. For RNA-seq, worms were stored in Trizol. For ChIP, worms were crosslinked in 2% formaldehyde for 30 minutes, followed by quenching in 125 mM glycine for 5 minutes, one wash with M9 and two washes with PBS, PMSF and protease inhibitors.

### Nuclei preparation for GRO-seq and Cut&Tag

Worms were resuspended in 2 mL of nuclei extraction buffer (20 mM HEPES–KOH, pH 7.9, 10 mM KCl, 0.1% Triton X-100, 20% Glycerol, 0.5 mM spermidine, protease inhibitors) and lysed by applying 100 strokes with a stainless-steel tissue grinder. Lysates were centrifuged at 100 g for 4 minutes at 4C to remove debris. Supernatants were transferred to 1.5 mL low binding microcentrifuge tubes and nuclei were pelleted by centrifuging at 1000 g for 4 minutes at 4C. Nuclei were washed 3 times with nuclei extraction buffer by repeated centrifugation at 1000 g for 4 minutes at 4C and resuspension in 1 mL of nuclei extraction buffer. For GRO-seq, nuclei were resuspended in 100 μL of freezing buffer (50 mM Tris-HCl pH 8, 5 mM MgCl_2_, 0.1 mM EDTA). For Cut&Tag, nuclei were resuspended in 500 μL of nuclei extraction buffer.

### GRO-seq

GRO-seq was performed as previously described^43^. Briefly, nuclear Run-On reactions were performed by incorporating 1 mM Bio-11UTP, followed by RNA extraction with Trizol and RNA fragmentation. Biotinylated nascent RNA was then purified using Dynabead^TM^ MyOne^TM^ Streptavidin C1 (Cat#65001). To prepare libraries, purified RNAs were first treated with T4 Polynucleotide Kinase (New England Biolabs) to repair the 5’-OH ends. This was followed by 3’ and 5’ adaptor ligation. Adaptor-ligated RNAs were then reverse transcribed using SuperScript IV Reverse Transcriptase (Thermo Fisher Scientific), following the manufacturer’s protocol with slight modifications: the reaction was incubated for 1 hour at 50°C and 10 minutes at 80°C. cDNA was PCR amplified using specific primers and the NEBNext Ultra II Q5 Master Mix 2x (New England Biolabs) for 15 cycles. Libraries were analyzed using the Agilent 2200 TapeStation System with high sensitivity D1000 screentapes and quantified with the Qubit Fluorometer High Sensitivity dsDNA assay kit (Thermo Fisher Scientific, Q32851). The multiplexed libraries were then sequenced on the Illumina NextSeq 2000 platform. Analysis of the sequencing data was performed as described in Quarato, et al 2021^43^. The scripts and workflows are available at https://gitlab.pasteur.fr/bli/bioinfo_utils. Two biological replicates were performed for each experiment.

### Cut&Tag

Cut&Tag was performed following the EpiCypher protocol with few modifications. Briefly, 100 μL of the nuclei resuspension generated as described above were used for binding to Convanavalin A beads (Epicypher 21-1401). Bead-bound nuclei were resuspended in 100 μL of Antibody150 buffer (20 mM HEPES pH 7.5, 150 mM NaCl, 0.01% digitonin, 2 mM EDTA, protease inhibitors, 0.5 mM spermidine) and 1 μg of primary antibody was added. Reactions were incubated overnight in a thermocycler shaker set up at 4 C with shaking cycles at 2000 rpm 15 seconds ON 45 seconds OFF. 1 μg of secondary antibody was added followed by tagmentation with Tn5 (Epicypher 15-1117). Resulting libraries were amplified with i5 and i7 primers using 13 PCR cycles and purified using AMPure beads. Libraries were analyzed using the Agilent 2200 TapeStation System with high sensitivity D1000 screentapes and quantified with the Qubit Fluorometer High Sensitivity dsDNA assay kit (Thermo Fisher Scientific, Q32851). Three biological replicates were performed for each experiment.

### Cut&Tag data processing and analysis

Reads were aligned with Bowtie2^44^ to the WBcel235 genome, with the following options: “—local--very-sensitive-local --dovetail --soft-clipped-unmapped-tlen”. Reads were additionally deduplicated using samtools and filtered to keep those with a single discovered alignment. Peaks were called using MACS2 with the following parameters: “callpeak-q 0.05-g ce --keep-dup all”. Peaks were further filtered to keep those with MACS2 FDR < 0.01 in merged reads from all conditions, and FDR < 0.05 in at least two individual replicates.

### RNA-seq

Total RNA was purified following Trizol manufacturer’s instructions after freeze-cracking samples five times. DNase treatment was performed using 2 μg of total RNA by treating with 2 U Turbo DNase (Ambion) at 37 C for 30 min followed by acid phenol extraction and ethanol precipitation. The Agilent 2200 TapeStation System was used to calculate the RIN indexes of RNA samples, and only samples with RIN > 8 were used for library preparation. For strand-specific RNA-seq library preparation, DNase-treated total RNA was used for ribosomal and mitochondrial rRNAs depletion following a custom RNAse-H-based method to degrade rRNAs based on complementary oligos, as described in Barucci et al., 2020^45^. Strand-specific RNA libraries were prepared using around 1μg of rRNA depleted RNAs with the NEBNext Ultra II Directional RNA Library Prep Kit for Illumina (E7760S). The size distribution of the RNA libraries was determined with Agilent 2200 TapeStation System using high sensitivity D1000 screentapes. Libraries were quantified using the Qubit Fluorometer High Sensitivity dsDNA assay kit (Thermo Fisher Scientific, Q32851). Two biological replicates were performed for each experiment.

### RNA-seq data processing and analysis

RNA-seq reads were aligned to the WBcel235 genome and ERCC92 spike-in sequences using STAR ^46^ and quantified by RSEM in parallel^47^ using the RSEM-STAR pipeline with the following additional options: ‘--calc-pme --calc-ci --estimate-rspd’. The distribution of spike-in counts across samples was used to estimate sample-specific normalization factors. Endogenous gene counts were then scaled using these factors to adjust for technical variation in library construction and sequencing efficiency. Further analysis was conducted using the Bioconductor DESeq2 package^48^ from gene-level values for rounded RSEM counts. Genes with at least 10 normalized counts in all replicates were considered for differential expression analysis. Differential expression between replicate groups was assessed using Wald Test, thresholding for significance at: FDR <= 0.05. Principal Component Analysis (PCA) and data visualization were made in R using’prcomp’ function and ggplot2 package^49^.

### ChIP-seq

Approximately 100 μL of pelleted L2 larvae were resuspended in FA buffer (150 mM NaCl, 50 mM HEPES/KOH pH 7.5, 1 mM EDTA, 1% Triton X-100, 0.1% sodium deoxycholate), supplemented with PMSF and protease inhibitors (Calbiochem, 539131), and lysed with 30 strokes of a glass homogenizer (Douncer). The lysates were sonicated for 15 minutes in 0.1% sarkosyl using a Bioruptor-pico. Protein concentration was measured using the Bradford assay (Biorad 500-0006), and 2 mg of protein extract were used for each ChIP (volume: 460 μL). A 5% aliquot was reserved as input DNA. The remaining protein extract was incubated with 5 μg of 8WG16 antibody (Millipore 05-952-I) at 4°C overnight with rotation. Next, 20 μL of Protein A and G Sepharose beads were washed three times with FA buffer and added to the immunoprecipitation reaction, which was rotated at 4°C for 2 hours. The beads were then washed with 1 mL of each of the following buffers: twice with FA buffer, once with FA-1 mM NaCl, once with FA-500 mM NaCl, once with TEL buffer (0.25 M LiCl, 1% NP-40, 1% sodium deoxycholate, 1 mM EDTA, 10 mM Tris-HCl pH 8.0), and twice with TE buffer. Immunoprecipitated DNA was eluted from the beads by incubating with ChIP elution buffer (1% SDS, 250 mM NaCl, 10 mM Tris pH 8.0, 1 mM EDTA) at 65°C for 30 minutes and then reverse-crosslinked at 65°C overnight in a water bath. For library preparation, half of the ChIP DNA and 30 ng of input DNA were ligated to Illumina TruSeq adapters and amplified by PCR. DNA fragments between 250 and 600 bp were gel-purified using the Qiagen gel extraction kit. Sequencing was performed using single-end 75 bp reads on the Illumina NextSeq 500 at the New York University Center for Genomics and Systems Biology. Two biological replicates with matching input samples were performed for each experiment.

### ChIP-seq data processing and analysis

Bowtie2 version 2.4.2 was used to align 75 bp single-end reads to the WS220 reference genome using default parameters^44^. BAM file sorting and indexing were performed with Samtools version 1.11^50^. BamCompare tool in Deeptools version 3.5.0 was used to normalize for the sequencing depth using CPM and create ChIP-Input coverage and ChIP/Input ratios with a bin size of 10 bp and 200 bp read extension^51^. Only reads with a minimum mapping quality of 20 were included, and mitochondrial DNA, PCR duplicates, and blacklisted genomic regions were excluded^52^. To generate average coverage data, ChIP-Input enrichment scores were averaged across 10 bp bins throughout the genome.

### FLAM-seq

FLAM-seq libraries were generated following previously established protocol^30^ (detailed version available at https://doi.org/10.21203/rs.2.10045/v1), starting with either 3 µg of total RNA from whole worms or 2 µg of total RNA isolated from worm nuclei. Each reaction included 4 µL of a 1:100 dilution ERCC RNA spike-in Mix 1 (Thermo Fisher). Polyadenylated RNA was enriched using the Stranded mRNA prep kit (Illumina), GI-tailed with reagents from the USB poly(A) Length Assay Kit (Thermo Fisher) and purified with 1.8x RNAClean XP beads (Beckmann Coulter). Reverse transcription was performed using the SMARTScribe Reverse Transcriptase kit (Takara Bio) with isoTSO and RT primer 1. Subsequently, cDNA was purified with 0.6x AMPure XP beads (Beckmann Coulter) and amplified by PCR using the Advantage 2 DNA Polymerase mix (Clontech). Amplified libraries were re-purified with 0.6x AMPure XP beads, followed by ligation to SMRTbell adapters, and sequencing on a PacBio Revio platform using a single SMRT Cell.

### FLAM-seq data processing and analysis

FLAM-seq datasets were processed using the FLAMAnalysis pipeline (https://github.com/rajewsky-lab/FLAMAnalysis) to determine poly(A) tail lengths and gene assignments. Reads were mapped using STARlong^46^ to *C.elegans* WBCel235 reference genome supplemented with ERCC spike-in reference sequences (Thermo Fisher). The resulting alignments were used to compute coverage and polyadenylation signal (PAS, AWUAAA) frequency in a window of 100 nucleotides around the WBCel235 annotated 3’ ends and the mapped 3’ ends respectively.

---

**Supplementary Figure 1.**
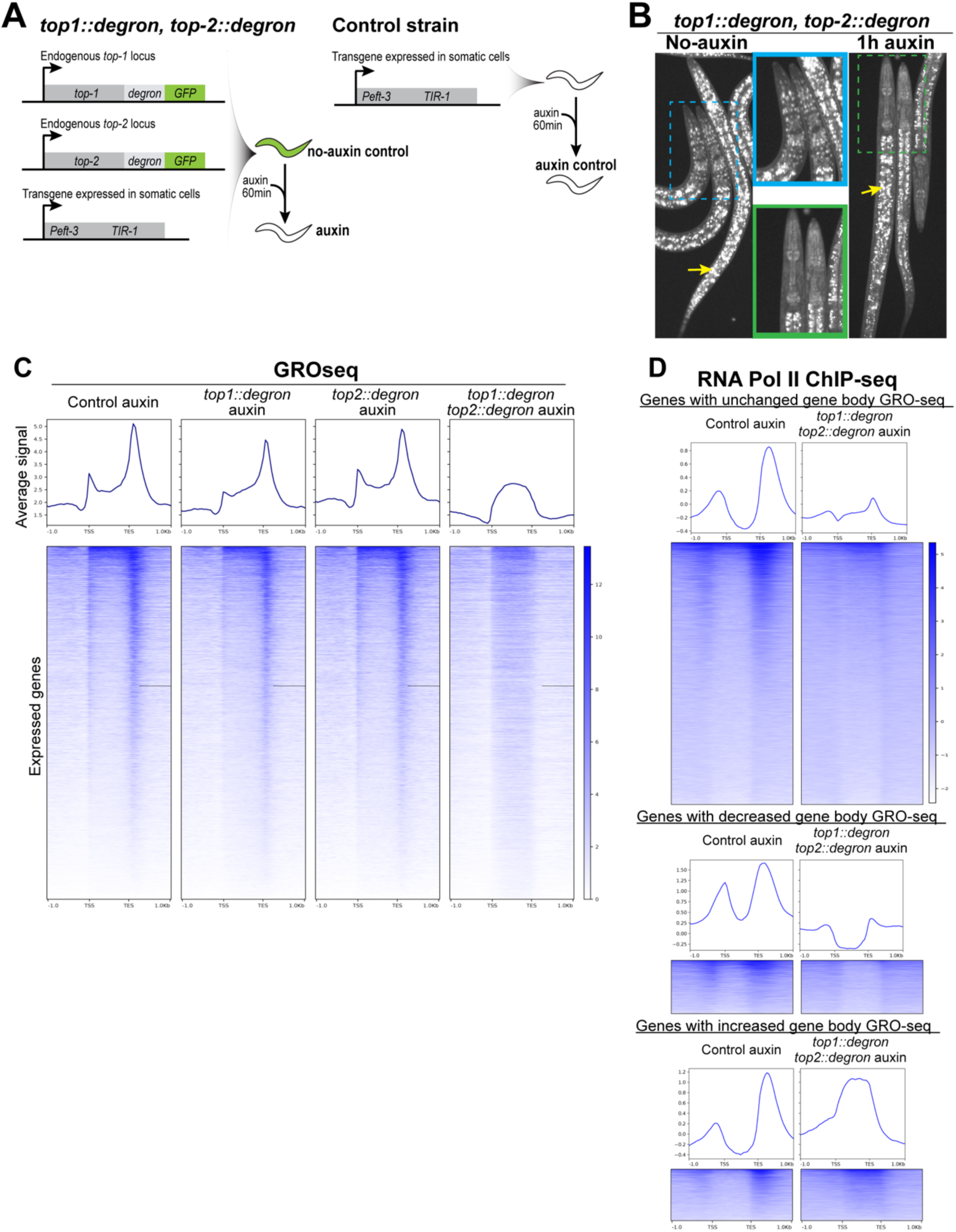
**A** Schematic representation of the worm strains used for the auxin-inducible degradation of TOP-1 and TOP-2. As controls, we used degron-tagged worms that were not treated with auxin (no-auxin control), and a strain without the degron tag but containing the TIR-1 transgene, that was treated with auxin (auxin control). **B** Images of *top-1::degron::GFP, top-2::degron::GFP* worms incubated for one hour in plates containing auxin. Yellow arrows indicate autofluorescent gut granules present throughout the intestine. **C** Heatmap showing GRO-seq signal across all expressed genes in *top-1::degron, top-2::degron and top-1::degron, top-2::degron* no-auxin and auxin conditions. Average plots are included at the top. **D** Heatmaps showing RNA Pol-II ChIP-seq scores across genes with unchanged, decreased and increased levels of GRO-seq signal within the gene body in the control auxin and *top-1::degron,top-2::degron* auxin conditions. Average plots are included at the top of the heatmaps.

**Supplementary Figure 2.**
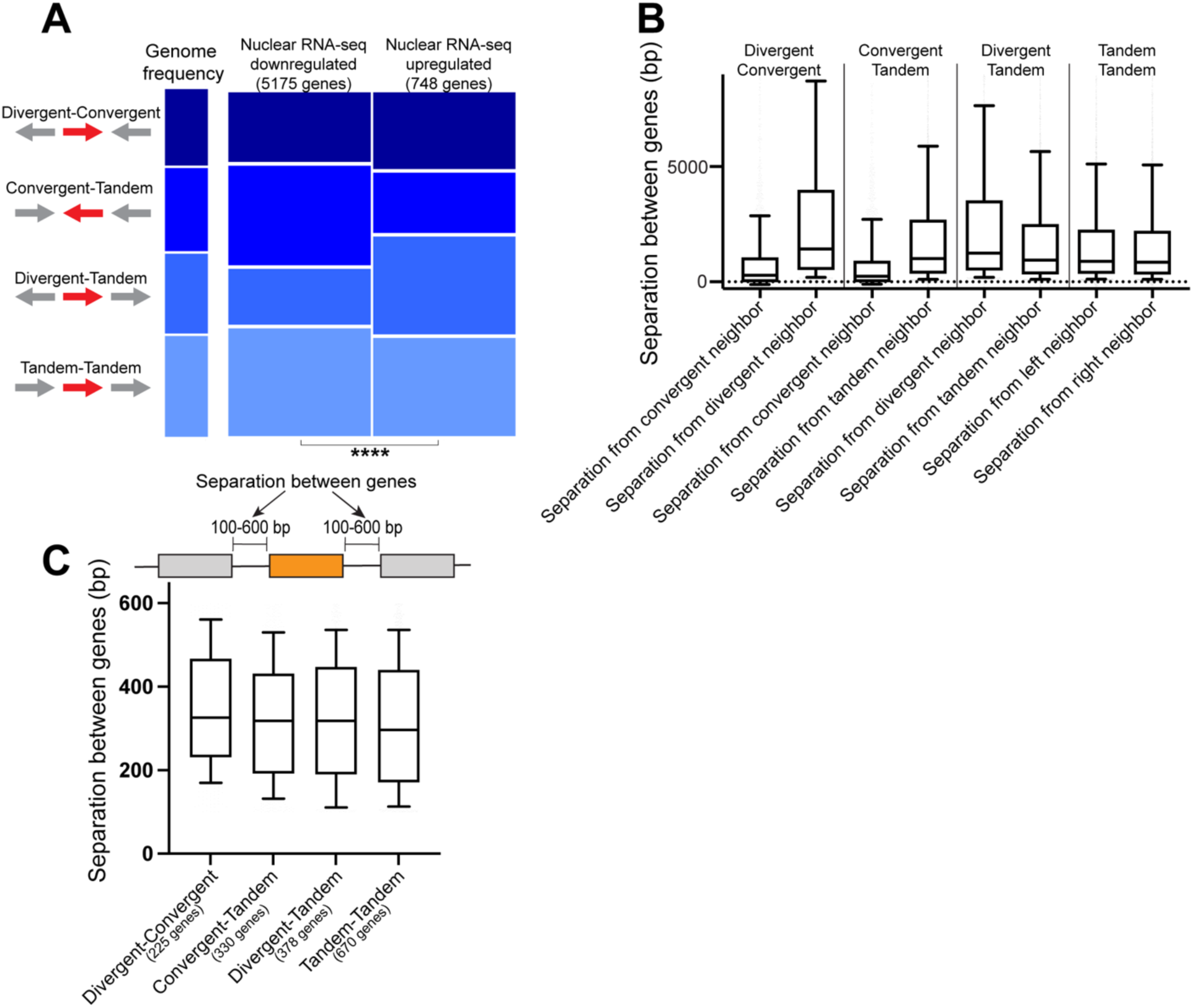
**A** Genes were classified based on the orientation of their upstream and downstream neighbors into four groups colored in different tonalities of blue: Divergent-Convergent, Convergent-Tandem, Divergent-Tandem and Tandem-Tandem. The height of the of the bars represents the proportion of genes belonging to each orientation group. This proportion is shown for the genome (left) and for genes found down and upregulated by nuclear RNA-seq upon topoisomerase depletion. **B** Distribution of the intergenic distance between genes neighbors is shown for genes grouped based on the orientation of their upstream and downstream neighbors. Boxplots display median (line), first, and third quartiles (box), and 90th/10th percentile values (whiskers). **C** Distribution of the intergenic distance between genes neighbors is shown for genes grouped based on the orientation of their upstream and downstream neighbors. Only genes with neighbors within a distance between 100-600 bp were included. Boxplots display median (line), first, and third quartiles (box), and 90th/10th percentile values (whiskers).

**Supplementary Figure 3.**
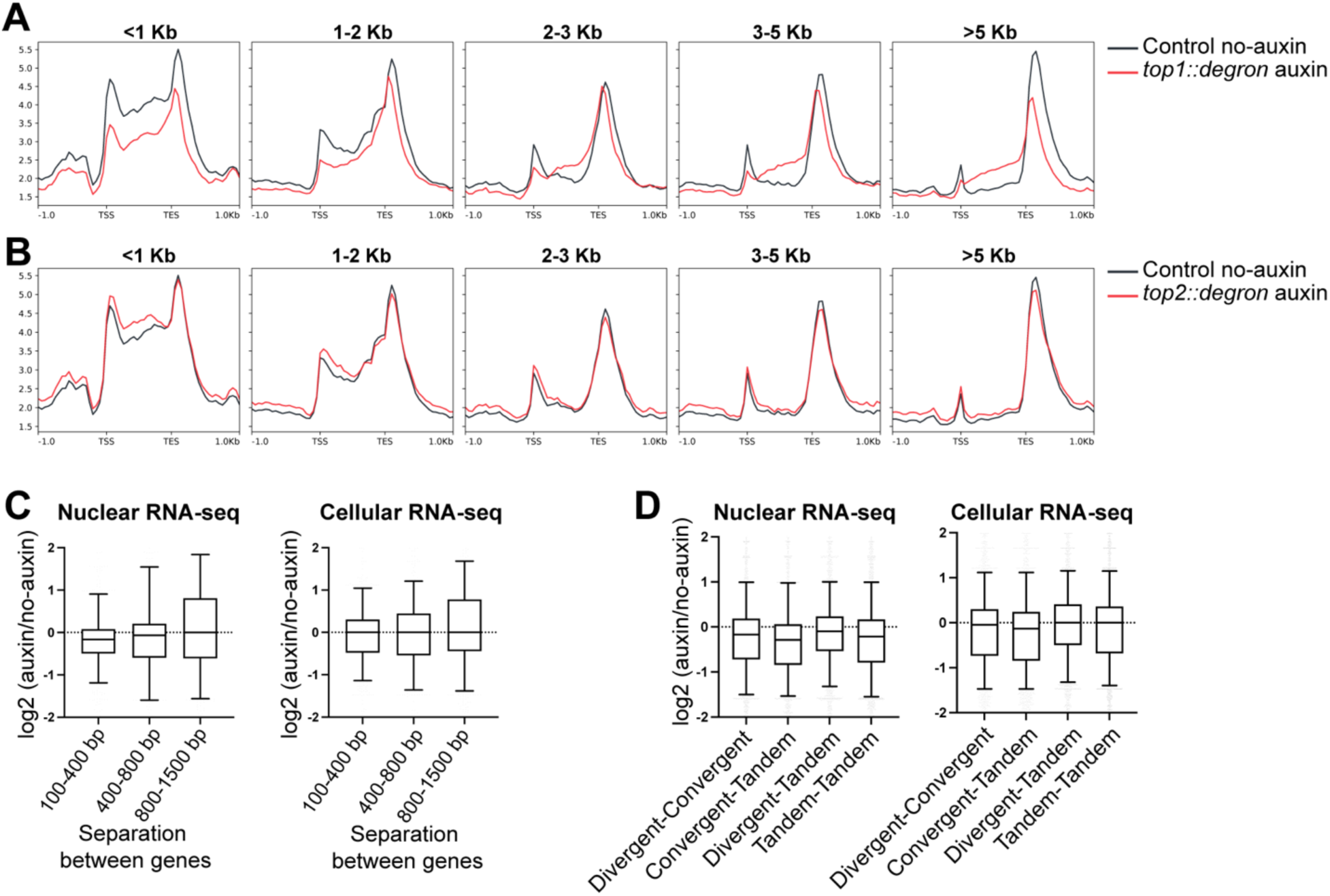
**A** Average GRO-seq scores in control (black) and TOP-1 depletion (red) are plotted across genes grouped based on their length into five categories: <1 Kb, 1-2 Kb, 2-3 Kb, 3-5 Kb and >5 Kb. **B** Average GRO-seq scores in control (black) and TOP-2 depletion (red) are plotted across genes grouped based on their length into five categories: <1 Kb, 1-2 Kb, 2-3 Kb, 3-5 Kb and >5 Kb. **C** Nuclear and cellular RNA-seq log2 fold changes between the auxin and no-auxin conditions are shown for genes grouped based on the distance separating them from their adjacent neighbors. Boxplots display median (line), first, and third quartiles (box), and 90th/10th percentile values (whiskers). **D** Nuclear and cellular RNA-seq log2 fold changes between the auxin and no-auxin conditions are shown for Divergent-Tandem, Tandem-Tandem, Divergent-Convergent and Convergent-Tandem genes. Boxplots display median (line), first, and third quartiles (box), and 90th/10th percentile values (whiskers).

**Supplementary Figure 4.**
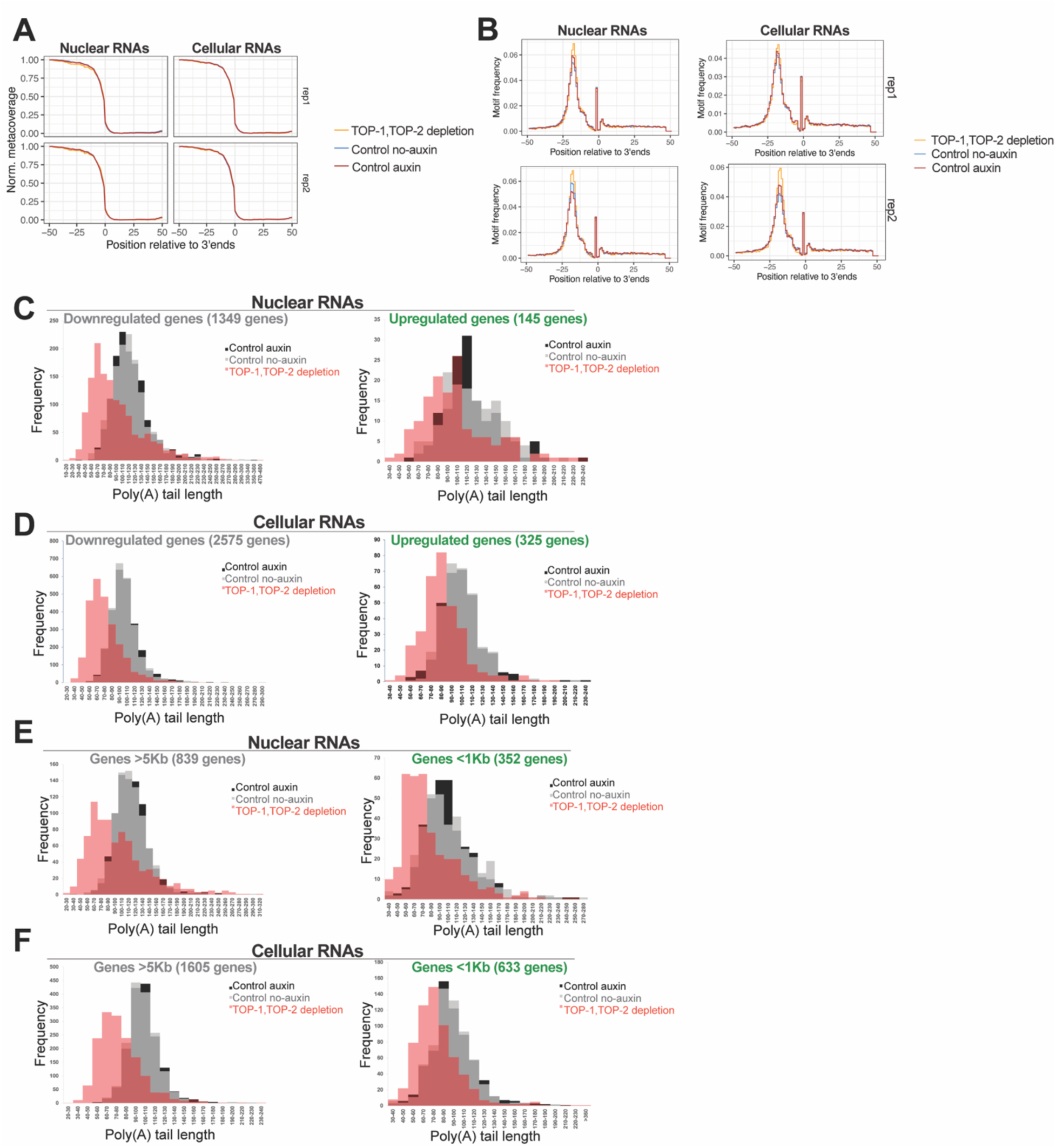
**A** FLAM-seq coverage in a 100 nt window around all transcripts’ 3’ ends for nuclear (left) and cellular (right) RNA in the controls (red and blue) and TOP-1/TOP-2 depletion (yellow). **B** Positional frequency around annotated 3’ ends of the polyadenylation motif A(A/U)UAAA in a 100 nt window around all transcripts’ 3’ ends for nuclear (left) and cellular (right) RNA in the controls (red and blue) and TOP-1/TOP-2 depletion (yellow). **C** Histograms of the distribution of poly(A) tail lengths of nuclear mRNAs in the controls (black and grey) and TOP-1/TOP-2 depletion (red) for genes found differentially expressed by nuclear RNA-seq upon topoisomerase depletion. **D** Histograms of the distribution of poly(A) tail lengths of cellular mRNAs in the controls and TOP-1/TOP-2 depletion for genes found differentially expressed by nuclear RNA-seq upon topoisomerase depletion. **E** Histograms of the distribution of poly(A) tail lengths of nuclear mRNAs in the controls and TOP-1/TOP-2 depletion for short (<1Kb) and long (>5Kb) genes. **F** Histograms of the distribution of poly(A) tail lengths of cellular mRNAs in the controls and TOP-1/TOP-2 depletion for short (<1Kb) and long (>5Kb) genes.

**Supplementary Figure 5.**
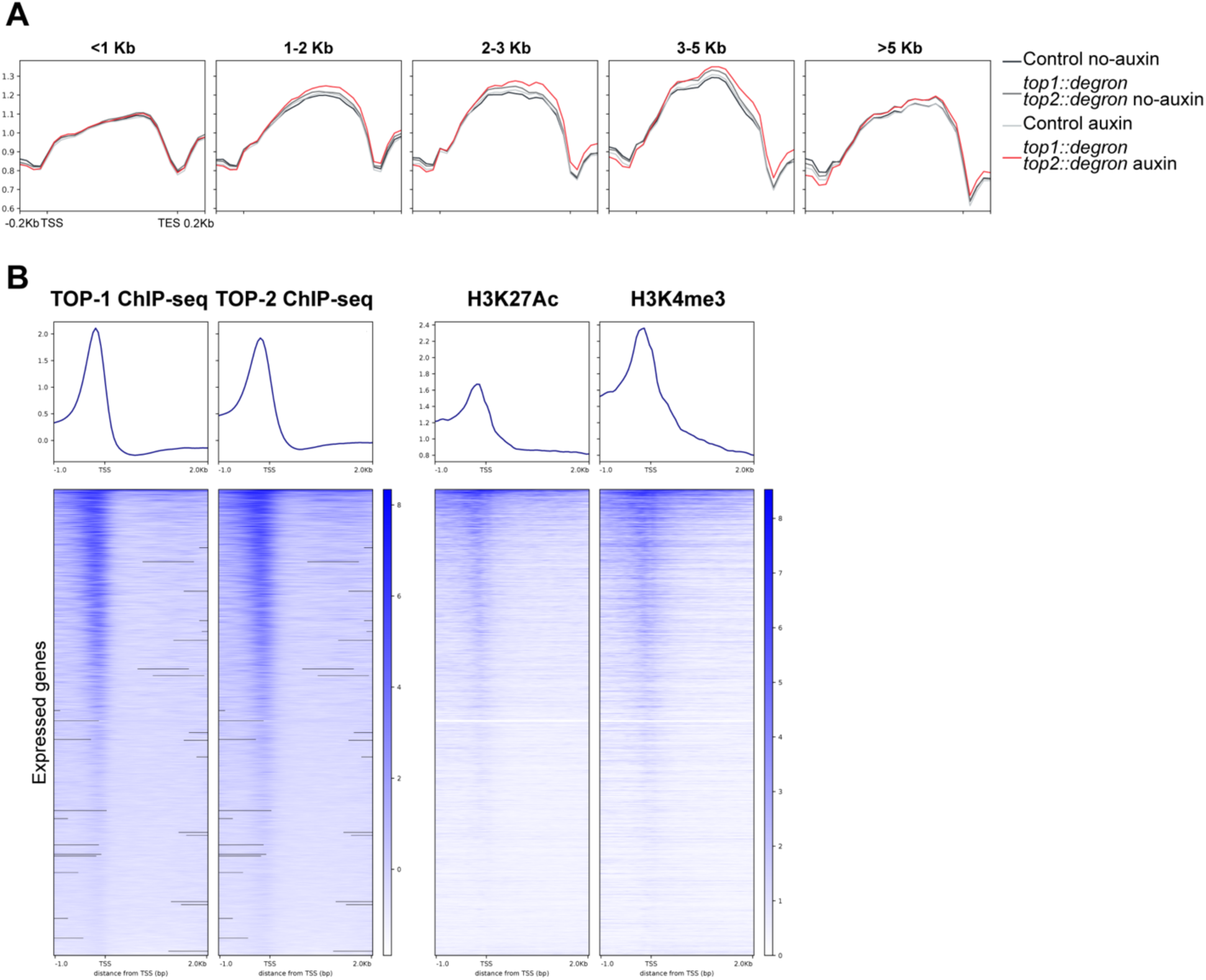
A. Average H3K36me3 scores in controls (black and gray) and TOP-1/TOP-2 depletion (red) are plotted across genes grouped based on their length into five categories: <1 Kb, 1-2 Kb, 2-3 Kb, 3-5 Kb and >5 Kb. **B** Heatmap showing TOP-1 and TOP-2 ChIP-seq scores (data from Morao et al, 2022), and H3K27Ac and H3K4me3 Cut & Tag control signal across promoters of expressed genes. Average plots controls are included at the top.

**Supplementary Figure 6.**
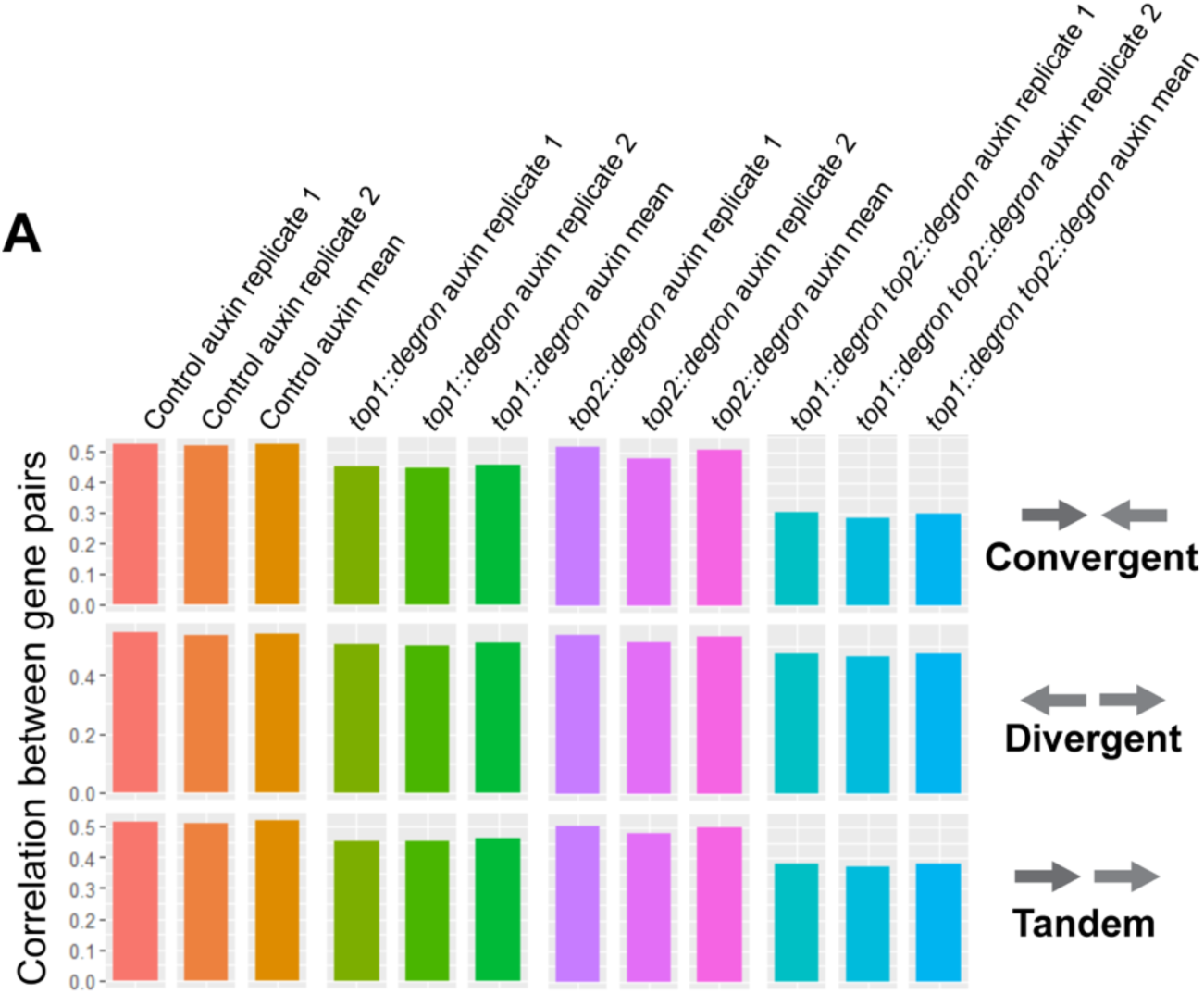
**A** Kendal correlations between GRO-seq TPM values of gene pairs with different orientations are shown for control, TOP-1 depletion, TOP-2 depletion and TOP-1/TOP-2 depletion. Values for replicates and average are included.

**Table S1.** List of the *C. elegans* strains used in this study.

| **strain name** | **short name** | **short description** | **strain genotype** | **Source** |
| --- | --- | --- | --- | --- |
| ERC83 | SS01 | *top-2::degron::GFP* at endogenous location in the CA1200 strain. | ers55[*top-2::degron::GFP*] II;<br>ieSi57 [eft-3p::*TIR1::mRuby::unc-54* 3'UTR + *Cbr-unc-119(+)*] II | Morao *et al*, 2022 |
| ERC84 | AM05 | *top-1::degron::GFP* at endogenous location in the CA1200 strain. | ers56[*top-1::degron::GFP*] I;<br>ieSi57 [eft-3p::*TIR1::mRuby::unc-54* 3'UTR + *Cbr-unc-119(+)*] II | Morao *et al*, 2022 |
|  |  | *top-1::degron::GFP* at endogenous location,<br>*top-2::degron::GFP* at endogenous location in the CA1200 strain. | ers56[*top-1::degron::GFP*] I;<br>ers55[*top-2::degron::GFP*] II;<br>ieSi57 [eft-3p::*TIR1::mRuby::unc-54* 3'UTR + *Cbr-unc-119(+)*] II | Morao *et al*, 2022 |
| CA1200 |  | Single copy transgene inserted into chromosome II (oxTi179) expressing modified *Arabidopsis thaliana* TIR1 tagged with mRuby in the soma | ieSi57 [eft-3p::*TIR1::mRuby::unc-54* 3'UTR + *Cbr-unc-119(+)*] II | CGC |

**Table S2.**
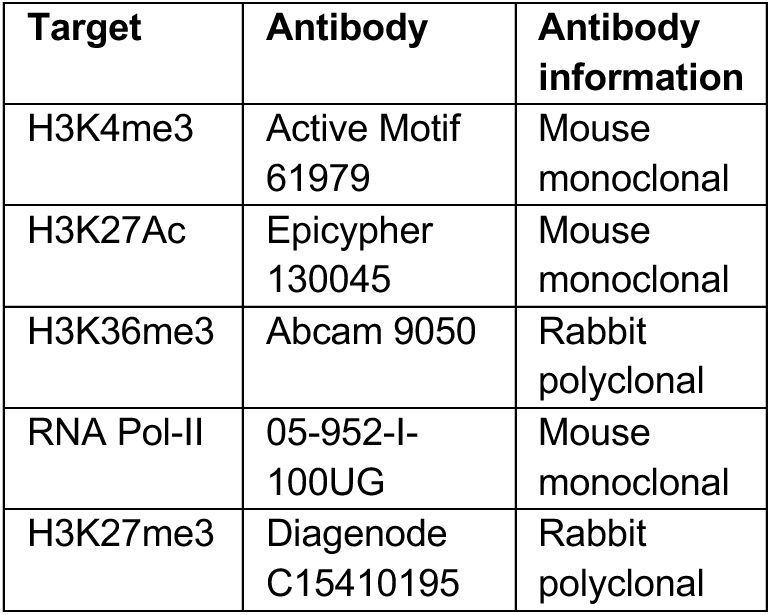
List of the antibodies used in this study.

## Notes

### Competing Interest Statement

The authors have declared no competing interest.

### Summary of Updates

Addition of new data and new authors. All figures have been revised.

